# Nine-residue low-complexity disordered peptide as a model system, an NMR/CD study

**DOI:** 10.1101/2023.02.16.528870

**Authors:** Liliya Vugmeyster, Aryana Rodgers, Kirsten Gwin, Dmitry Ostrovsky, Serge L. Smirnov

**Affiliations:** Department of Chemistry, University of Colorado Denver, Denver CO USA 80204; Department of Mathematics, University of Colorado Denver, Denver CO USA 80204; Department of Chemistry, Western Washington University, Bellingham, WA 98225

**Keywords:** low-complexity peptides, peptide NMR, disordered regions, NMR relaxation of peptides

## Abstract

Disordered proteins and protein segments can be crucial for biological function. In this work we present a detailed biophysical characterization of the low-complexity nine-residue peptide with the sequence GGKGMGFGL. Based on proton solution NMR chemical shifts, circular dichroism measurements, as well as the analysis of concentration dependence of NMR linewidth, proton longitudinal relaxation times, hydrogen-deuterium exchange measurements, and ^15^N rotating frame NMR relaxation measurements, we conclude that the peptide is fully disordered and monomeric in solution. The peptide will serve as a model system for future structural and dynamics studies of biologically relevant disordered peptides in solution and solid states.

## Introduction

Disordered proteins and disordered regions are recognized to play an important part in various biological phenomena such as ligand binding, signaling events, liquid-liquid phase separation, and many others.^[1-2]^ In this work we study a nine-residue low-complexity^[3-4]^ peptide (RC9) designed as a model unstructured unit of a disordered system, with the sequence: GGKGMGFGL. On one hand, the sequence is designed to have high solubility due to the presence of a polar sidechain, and, on the other hand, span a selection of methyl and aromatic groups. The latter will allow for follow up studies of mobile side-chains in solution and solid states.^[5]^ The peptide can be made synthetically.

Here, we focus on the assessment of the degree of disorder using conventional homonuclear solution NMR resonance assignments supplemented with the natural abundance C-alpha backbone chemical shifts, as well as circular dichroism (CD) measurements. Further, we employ one dimensional ^1^H NMR hydrogen-deuterium(H/D) exchange measurements to quantify the extent of exchange with the solvent and, thus, further confirm the extent of disorder. To test for the monomeric state, we examine the linewidth at two different protein concentrations and scan for the potential presence of non-sequential NOE’s. ^15^N rotating frame *R*_1*ρ*_ NMR relaxation experiments on the selectively labeled ^15^N amide of the F7 residues are performed to further exclude monomer-dimer exchange or any slow time scales motions of the backbone on time scale within the sensitivity range of these measurements.

## Results

### Solution NMR resonance assignments indicate a disordered structure

Amide proton (^1^H_N_) NMR chemical shift values can be strong indicators of random coil or folded backbone in protein samples.^[6]^ Here, we utilized ^1^H 2D NOESY (300 ms mixing time), ROESY (300 ms) and TOCSY (45 ms and 90 ms) measurements recorded at peptide concentration of 2 mM and 285 K to perform backbone and side chain proton chemical shift assignments using the established procedure.^[7]^ An acidic solution condition of pH 2 was selected to maximize the NMR signal intensities of amide protons and to ensure the protonated state of the ionizable side chain groups. At neutral pH and at 285 K all amide proton resonances were not visible on 1D proton spectrum, indicating fast exchange with the solvent. Figure 1A shows representative ^1^Hα-^1^H_N_ and ^1^Hα-side chain ROESY and TOCSY signals for the non-glycine spin systems (K3, M5, F7 and L9). In addition, Figure 1B presents TOCSY ^1^H_N_-^1^Hα resonances for G4, G6 and G8 as well as sequential ROESY ^1^Hα(i)-^1^H_N_(i+1) signals for the bulk of the sequence. Most of the resonances originating from the non-glycine side chain spin systems were reliably and unambiguously identified (Table S1). The ^13^Cα chemical shift values for all the non-glycine residues (K3, M5, F7 and L9) were assigned from the combined use of ^1^Hα – ^13^Cα resonances in the natural abundance ^13^C-HSQC and the ^1^Hα – ^1^H_N_ resonances in ^1^H 2D TOCSY (45 ms) spectra (Figure S1). All the backbone amide chemical shift values for sequence positions K3 through L9 are within 8.0 – 8.6 ppm range, which is consistent with an unfolded, random coil conformation. The ^1^H_N_ chemical shift values of the three internal glycine residues (G4, G6 and G8) were unambiguously identified through the sequential ROESY (300 ms) signals with the ^1^Hα resonances of the previous residues (K3, M5 and F7 respectively) as shown in Figure1B (top panel, signals A, C and E). The backbone amide ^1^H_N_ and ^1^Hα resonances for the G1 residues were assigned from their low-intensity TOCSY cross-peak (Supplementary Figure S2. This G1 TOCSY off-resonance signal corresponds to one of the two broad peaks in 1D ^1^H spectra originating from NH_3_^+^ groups, while the other one originates from the K3 side chain (Figure 3). The ^1^H_N_ chemical shift value of the G2 residue can be inferred from the 1D ^1^H signal patters (Figure 3). The most downfield peak has a distinct multiplet structure and at these conditions most likely corresponds to an overlap of resonances of the G2 and G4 amide protons. Each ^1^H_N_ signal originating from an internal glycine residue (e.g., G8, Figure 3) is split into a 1:2:1 pattern due to an intra-residual J-coupling with two ^1^Hα nuclei. The signals originating from the G2 and G4 amide protons are likely shifted with respect to each other by the value corresponding to 2*x*J-coupling (J = 5.75 Hz), which explains the observed 1:2:2:2:1 pattern (Figure 3). The assignments were also validated using the toolset in POKY.^[8]^ Importantly, no resonance patterns indicating an internal fold, a specific secondary structure or a dimer/oligomer state of the sample were detected.

**Figure 1.**
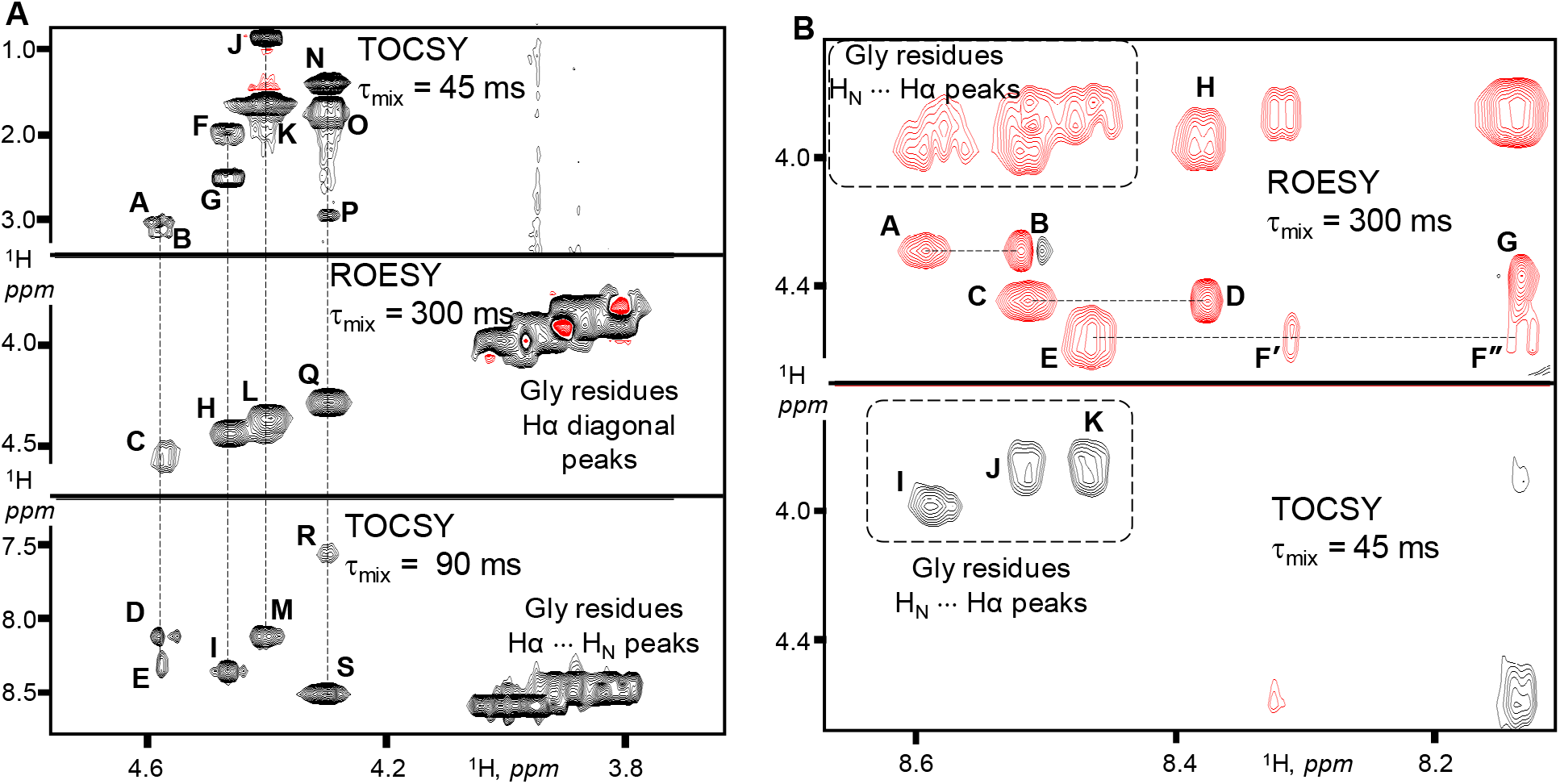
2D proton NMR spectra of RC9 peptide used for NMR chemical shift assignments, collected at 11.7 T field strength, 12 °C, 2 mM concentration and pH 2. Panel **A**: ^1^H TOCSY (off-diagonal signals) and ^1^H ROESY (diagonal signals, black for positive values) representing K3, M5, F7, and L9 side-chain spin systems. Signals **A**-**E**, F7 spin-system: **A** (Hα – Hβ3), **B** (Hα – Hβ2), **C** (Hα diagonal), **D**/**E** (Hα – H_N_, ^1^H split due to ^15^N). Signals **F**-**I**, M5 spin-system: **F** (Hα – Hβ3/Hβ2 overlap), **G** (Hα – Hγ3/Hγ2), **H** (Hα diagonal), **I** (Hα – H_N_). Signals **J**-**M**, L9 spin-system: **J** (Hα - Hδ), **K** (Hα - Hγ), **L** (Hα diagonal), **M** (Hα – H_N_). Signals **N**-**S**, K3 spin-system: **N** (Hα – Hγ3/Hγ2), **O** (Hα – Hβ3/Hβ2), **P** (Hα – Hε), **Q** (Hα diagonal), **R** (Hα – Hζ), **S** (Hα – H_N_). Panel **B**: Sequential and intra-residual H_N_-Hα connectivities represented by off-diagonal ^1^H ROESY (red for negative values) and ^1^H TOCSY signals. ROESY: **A** (G4_H_N_ – K3_Hα), **B** (K3_H_N_ – K3_Hα); **C** (G6_H_N_ – M5_Hα), **D** (M5_H_N_ – M5_Hα); **E** (G8_H_N_ – F7_Hα), **F*’*** / **F*”*** (F7_H_N_ – F7_Hα), F7 backbone has an ^15^N label, thus the observed **F’** / **F”** ∼90 Hz split; **G** (L9_H_N_ – L9_Hα); **H** (M5_H_N_ – G4_Hα); TOCSY glycine H_N_ – Hα signals: **I** (G4), **J** (G6), **K** (G8).

**Figure 2.**
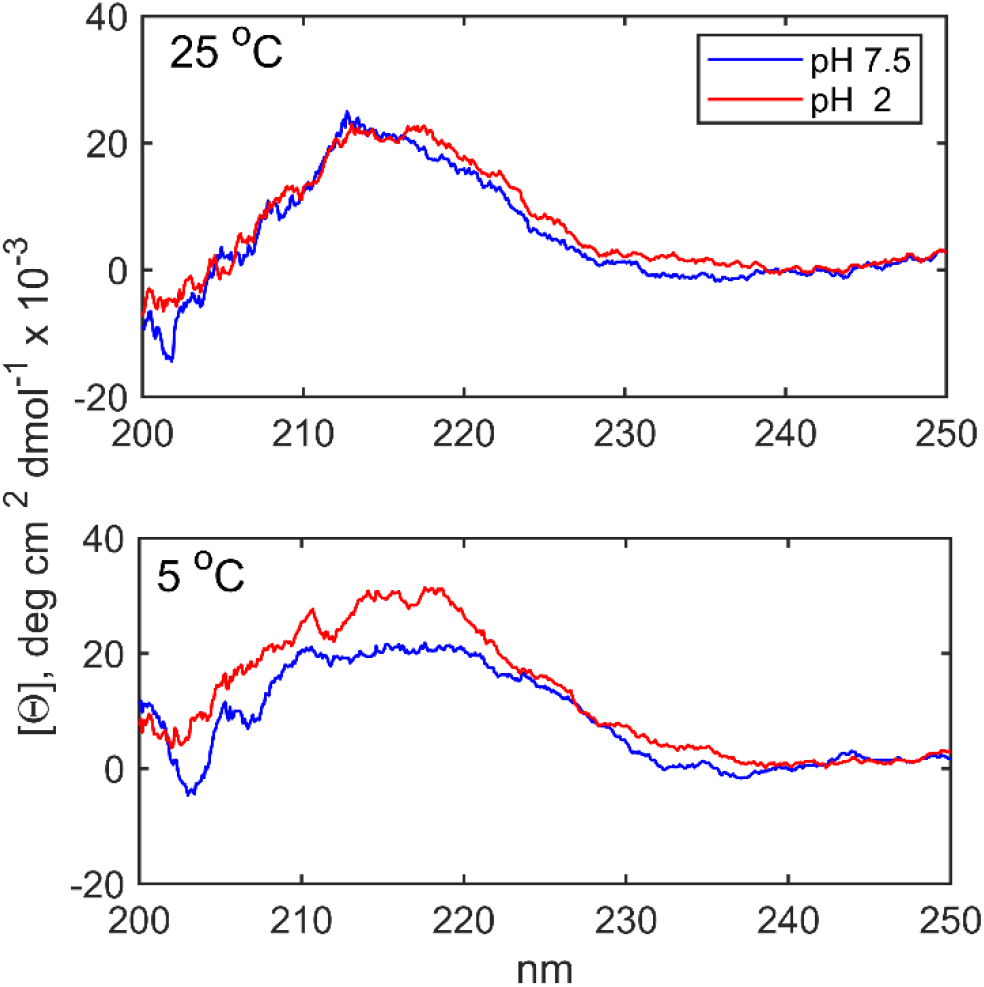
CD spectra of RC9 at 25 μM concentration recorded using Jasco-1500 spectropolarimeter with 1 cm pathlength quartz cuvettes.

**Figure 3.**
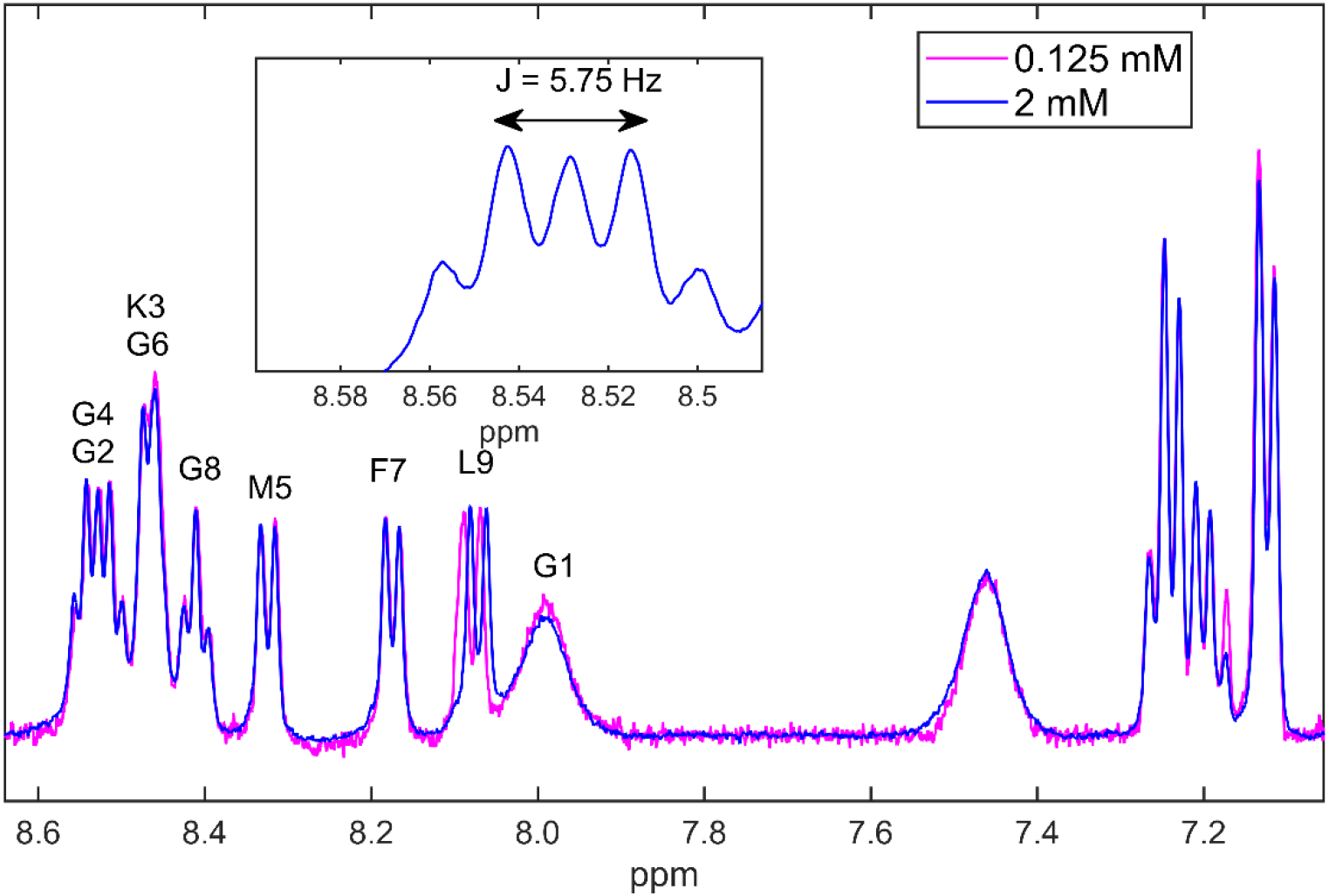
A scaled overlay of 1D proton NMR spectra of RC9 and 2 mM (blue) and 0.125 mM (magenta) concentrations at pH 2 and 7°C. The spectra were recorded at 9.4 T magnetic field strength using the broadband probe, The number of scans was 32 and 632 for the 2 mM and 0.125 mM concentrations, respectively. The assignments of the backbone amide region are shown explicitly. The insert on the top shows the distinct J-split pattern originating due to the shift between G2 and G4 ^1^H_N_ sites by 2J (2 × 5.75 Hz = 11.5 Hz). The backbone amide (the broad peak at ∼8.0 ppm) and Hα resonances for G1 were assigned from their TOCSY cross-peak (Figure S2). The second broad signal between 7.4 and 7.6 ppm originates from the side chain NH_3_^+^ group of K3.

### Circular Dichroism measurements further support the absence of secondary structure

CD measurements were performed in the 200 to 250 nm wavelength range at a protein concentration of 25 μM at two temperatures of 5 and 25 °C and at two pH values of 2 and 7.5 (Figure 2). All the CD spectra are consistent with a random coil state of the sample and do not detect any sizable presence of the secondary structure.^[9]^ Thus, the results of the CD measurements are consistent with the NMR assignments analyses, confirming the disordered structure of the backbone.

### Concentration dependence of 1D proton NMR spectral linewidth

The changes of the linewidth upon changes in concentration can report on presence of dimers and oligomers.^[6]^ We have taken 1D proton NMR spectrum at 2 and 0.125 mM concentrations, and their overlay in the 7.0 to 8.6 ppm chemical shift range is shown in Figure 3. Based on the expected scaling and taking into account the difference in the number of scans for the two concentrations, the expected peak intensity scaling factor was 0.81, and the experimental values ranged from 0.8 to 0.84. Thus, no dimer presence is detected. There is a slight apparent chemical shift change at the L9 site of 0.00721 ppm in the downfield direction (∼3.60 Hz) for the diluted peptide. We have also measured proton *T*_1_ relaxation times, presented in Figure S3. In the aromatic region there is no difference between the *T*_1_ values at the two concentrations, again excluding the presence of the dimer. In the amide region, the diluted sample has a tendency to have somewhat lower *T*_1_ values compared to the concentrated sample, suggesting slight differences in the fast backbone rearrangements, perhaps due to differences in solution viscosities.

### ^15^N R_1*ρ*_ NMR relaxation experiments provide further confirmation for the absence of monomer-dimer exchange

^15^N *R*_1ρ_ NMR relaxation measurements probe the relaxation rate as a function of effective spin-locking field. The experiments can be carried out on or off resonance,^[10]^ the latter allowing for a higher effective spin-locking field without additional RF heating effects. To align magnetization along the effective tilted axis in the off-resonance case, adiabatic ramps have to be employed.^[11]^ The presence of relaxation dispersion (i.e., the decrease of the relaxation rate with the increase of effective spin-lock field) indicates the presence of dynamics on μs-ms time scale, which can originate either from internal molecular motions or oligomerization.^[12]^ We have conducted such measurements for the selectively ^15^N-labeled amide backbone site of the F7 residue using two spin-locking field amplitude values of 1.5 kHz and 2.5 and off resonance offset values in −2 to +2 kHz range. No dispersion was observed in the relaxation rates and all rates ranged from 1.11 to 1.18 s^-1^. Representative data are shown in Figure S4. The lack of the dispersion indicates no dimerization is happening on the slow time scale and the backbone does not undergo rearrangements on the μs to ms time scale.

### Hydrogen-deuterium exchange measurements further tests for the degree of disorder

To further probe the extent of disorder in this peptide and quantify the exchange of protons with solvent, we have conducted H/D exchange measurement using 1D proton NMR. Due to anticipated fast rates the exchange was measured at 7 °C and pH 2. Upon dissolution of the peptide in D_2_O on ice, the first time point was collected at 12 min. The rate constants *k*_H/D_ obtained for all the backbone amide peaks, some of which overlap, are shown in Figure 4. The exchange was near completion for all sites after two hours. The raw data and fits are shown in Figure S5.

**Figure 4.**
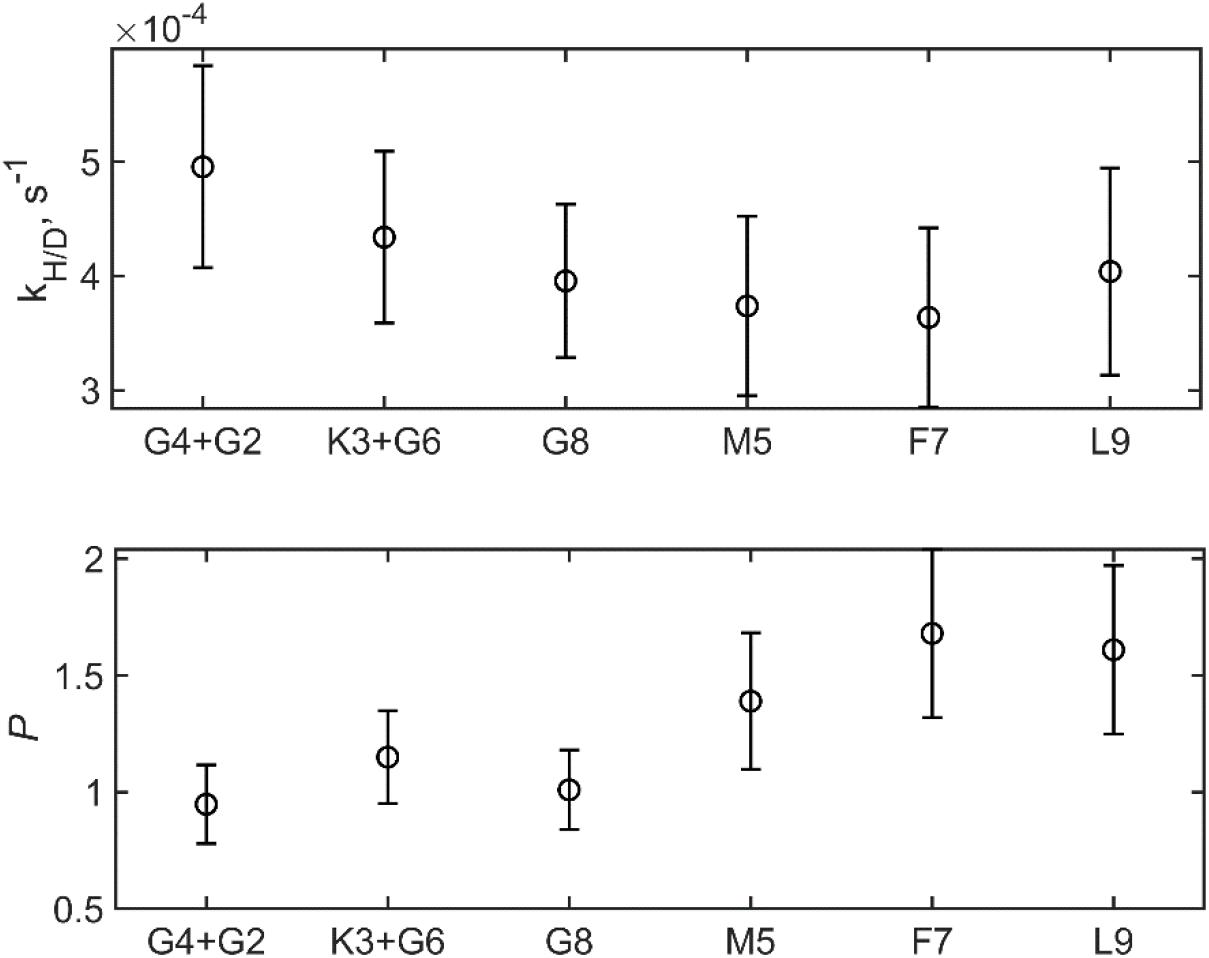
Top panel: Backbone amide protons *k*_H/D_ rate constants measured by 1D proton NMR at 7 °C, pH 2 at 9.4 T. Bottom panel: protection factors calculated as described in the text.

There is little variability in the resulting rate constants themselves. To determine the so-called protection factor *P*, one needs to divide the experimental rate by their corresponding random coil values. We have used the method by Bai et al. to obtain the random-coil values, ^[13]^ which is based on oligopeptide models with inclusion of inductive and steric effects, as well as the correction for the deuterium isotope effect on the pH value. For the overlapping peaks, the average random-coil values were used. In the absence of protection from the solvent the *P* is 1. Within the error bars all values are either 1 or very close to 1. M5, F7, and L9 sites display a tendency to have somewhat higher protection factors.

## Discussion and Conclusion

The biophysical evidence presented above points to RC9 remaining fully disordered and monomeric in solution at concentrations up to 2 mM. We plan to employ this peptide as a model system in future studies of low complexity disordered proteins in solution and solid states. At the same time, similar primary sequences or patterns can be found in naturally occurring intrinsically disordered regions (IDRs). For example, a search for the pattern *x*(R/K)*x*M*x*F*x*(P/L/A), where *x* is any residue of the same type, can be identified as the oligopeptide SRSMSFSP belonging to the intrinsically disordered region (IDR linker) within plant villin 4 from *A. thaliana* (NCBI accession NM_001203940). This sizable internal linker, spanning 192 residues, has a distinct charge-partitioning pattern: the 144-residue N-terminal basic segment with the isoelectric point is at 11.5 and the 48-residue C-terminal acidic one with the isoelectric point 4.1.^[14]^ The region has been shown to have a novel capacity to bind/bundle actin filaments.^[15]^ Notably, the closest match to the basic RC9 oligopeptide is basic (SRSMSFSP) and it is positioned within the basic part of the IDR in the villin 4 linker. Thus, identification of similar oligopeptide patterns within naturally occurring biologically relevant IDRs and investigation of the dynamics and structure of these shorter segments increases our understanding of the functional mechanisms of the larger IDRs.

## Materials and Methods

### Peptide preparation

The RC9 peptides with the sequence GGKGMGFGL was prepared using solid-state peptide synthesis with Fluorenylmethyloxycarbonyl (FMOC)-based chemistry by Life Technologies, Inc and purified by reversed-phase high pressure liquid chromatography (HPLC) using the C18 column and water/acetonitrile buffer system with trifluoroacetic acid as the co-factor. The purity of the peptide at the level of 98% was confirmed by analytical HPLC using the 0.5% acetonitrile/min gradient, and matrix-assisted laser desorption/ionization mass spectroscopy. FMOC-Met-^13^CH_3_, FMOC-Phe-^15^N, and FMOC-Leu-D_3_ were purchased from Cambridge Isotope laboratories.

The resulting peptide used in the chemical shift assignments studies had several isotopic labels, with the ^15^N label at the backbone amide site of F7 the only one relevant for this study. The other labels were at the methyl groups of M5(^13^C) and L9(D_3_). The second peptide, used in the concentration-dependence, longitudinal proton NMR relaxation and the H/D exchange measurements, did not have the ^15^N label at the backbone F7 site. For all NMR measurements except the H/D exchange studies the peptides were dissolved in 90% H_2_O/10% D_2_O containing 0.1% sodium azide and pH adjusted using HCl/KOH solutions.

### NMR measurements

Solution NMR assignments experiments (NOESY, ROESY, TOCSY) as well as on and off-resonance ^15^N *R*_1ρ_ measurements were performed at 12 °C, pH 2, and 2 mM concentration with the 90% H_2_0/10%D_2_O/0.1% NaN_3_ solution mixture as the solvent. These measurements were conducted using the Bruker Avance III HD NMR spectrometer operating at 11.7 Tesla (500 MHz ^1^H frequency) equipped with the triple-resonance TCI cryo-probe at Boston University School of Medicine. The ^1^H and ^13^C resonance assignments for RC9 are now available as Biological Magnetic Resonance Bank (BMRB) entry 51754.

^15^N *R*_1ρ_ NMR relaxation measurements measurement were performed in a one-dimensional fashion using the pulse sequence described in Mulder et al.^[11]^ The spin-lock field strength applied at the ^15^N site of F7 was either 1.5 or 2.5 kHz and the values of the off-resonance offsets were of 0, ±1 kHz, ±2kHz. Nine relaxation delays were utilized in the 1 ms to 188 ms range. The data reported represents the average of the two peaks split by the H_α_ scalar coupling. The decays were fitted to the mono-exponential function and the errors determined by covariance matrix method.^[16]^

The H/D exchange measurements, concentration dependence of the linewidth, and the ^1^H longitudinal times measurements at different concentrations were performed using Bruker 400 MHz spectrometer equipped with the broad-band double-resonance probe (CU Denver) at 7 °C and pH 2. No isotopic labels at the backbone were introduced for the peptide used in these measurements. The H/D exchange measurements were performed at 0.9 mM peptide and conducted in a 1D fashion using the watergate pulse sequence^[17]^ for water suppression. The first time point was 12.5 min counting from the time point of dissolving in D_2_O. The last collection point was at 128 min. The number of scans varied from 16 for the shortest times to 64 for the longest times. The data reported represents the average of the peaks split by the H_α_ scalar coupling.

The spectra were base-line corrected and the resulting intensities were fitted to a mono-exponential function with a baseline,^[18]^ with the errors determined from the covariance matrix method. The random coil values were calculated using the Sphere program.^[13]^

The concentration dependence of proton amide linewidths was taken at 2 mM and 0.125 mM peptide concentrations (i.e, dilution by a factor of 16) using the 90% H_2_0/10%D_2_O/0.1% sodium azide solution as the solvent. The corresponding ^1^H NMR spectra were collected using the watergate one-dimensional experiments. The spectral width was 6493.51 Hz, and the number of scans was 32 for the 2 mM and 632 for the 0.125 mM peptide concentration, respectively. 16384 complex FID points were collected in both cases. The spectra were processed with the 0.3 Hz line broadening function.

### CD measurements

CD experiments were conducted using Jasco-1500 spectropolarimeter with the use of 1 cm pathlength quartz cuvettes. The peptide concentration was 25 μM in milliQ water with 0.1% NaN_3_ added, and this solvent used as a reference. Spectra were averaged over three scans using scanning speed of 100 nm/min and 3 nm bandwidth.

## Acknowledgements

We thank Stowe Shawn for assistance with the CD measurements, Andrea Lopez for assistance with POKY software, and C.J. McKnight for access to the 500 MHz spectrometer at Boston University School of Medicine, supported by the NIH grant S10OD011941. The work was also supported by 1R15-GM111681 to L.V and NSF-MRI instrumentation grant CHE-1726947.

## Notes

### Competing Interest Statement

The authors have declared no competing interest.

### Summary of Updates

Fig-1, Table-S1, Fig-S1, S2 (minor differences in chemical shift referencing)

## References

[1] R. B. Berlow, H. J. Dyson, P. E. Wright, J. Mol. Biol. 2018, 430, 2309–2320.

[2] P. E. Wright, H. J. Dyson, Nat. Rev. Mol. Cell. Bio. 2015, 16, 18–29.

[3] A. C. Murthy, G. L. Dignon, Y. Kan, G. H. Zerze, S. H. Parekh, J. Mittal, N. L. Fawzi, Nat. Struct. Mol. Biol. 2019, 26, 637–648.

[4] A. C. Murthy, N. L. Fawzi, J. Biol. Chem. 2020, 295, 2375–2384.

[5] C. T. Mant, B. Tripet, R. S. Hodges, J. Chromatogr. A 2003, 1009, 45–59.

[6] J. Cavanagh, W. J. Fairborther, A. G. P. Palmer, N. J. Skelton, M. Rance, Protein NMR Spectroscopy. Principles and Practice, 2nd ed., Elsevier Academic Press, 2006.

[7] K. Wüthrich, G. Wider, G. Wagner, W. Braun, J. Mol. Biol. 1982, 155, 311–319.

[8] M. Rahimi, Y. Lee, H. Nguyen, A. Chiu, W. Lee, J. Magn. Reson. 2022, 339, 107214.

[9] S. M. Kelly, T. J. Jess, N. C. Price, Biochim. Biophys. Acta 2005, 1751, 119–139.

[10] A. G. Palmer, 3rd, J. Magn. Reson. 2014, 241, 3–17.

[11] F. A. A. Mulder, R. A. de Graaf, R. Kaptein, R. Boelens, J. Magn. Reson. 1998, 131, 351–357.

[12] A. G. Palmer, C. D. Kroenke, J. P. Loria, Methods Enzymol. 2001, 339, 204–238.

[13] Y. Bai, J. S. Milne, L. Mayne, S. W. Englander, Proteins 1993, 17, 75–86.

[14] K. V. Boyko, E. A. Rosenkranz, D. M. Smith, H. L. Miears, M. Oueld Es Cheikh, M. Z. Lund, J. C. Young, P. N. Reardon, M. Okon, S. L. Smirnov, J. M. Antos, PLoS One 2021, 16, e0258531.

[15] M. Zou, H. Ren, J. Li, Plant Physiol. 2019, 181, 161–178.

[16] J. Shao, Mathematical Statistics, Springer, New York, 2003.

[17] M. Piotto, V. Saudek, V. Sklenár, J. Biomol. NMR 1992, 2, 661–665.

[18] L. Vugmeyster, B. Kuhlman, D. P. Raleigh, Protein Sci. 1998, 7, 1994–1997.

