## Supporting Information for "Nine-residue low-complexity disordered peptide as a model system, an NMR/CD study"

**Table S1.** Summary of NMR chemical shift assignments for the RC9 peptide with the sequence GGKGMGFGL. Chemical shift values and precision are in *ppm*.

| Residue | Atom | Chemical Shift | Precision |
|---|---|---|---|
| **GLY 1** | HN | 8.072 | 0.01 |
| | HA2/HA3 | 3.891 | 0.02 |
| **LYS 3** | HA | 4.301 | 0.005 |
| | HB2 | 1.808 | 0.02 |
| | HB3 | 1.726 | 0.02 |
| | HD2/HD3 | 1.649 | 0.005 |
| | HE2/HE3 | 2.979 | 0.005 |
| | HG2/HG3 | 1.417 | 0.005 |
| | HN | 8.509 | 0.005 |
| | HZ | 7.542 | 0.01 |
| | CA | 56.46 | 0.05 |
| **GLY 4** | HN | 8.589 | 0.005 |
| | HA2/HA3 | 3.999 | 0.02 |
| **MET 5** | HA | 4.464 | 0.005 |
| | HB2 | 1.966 | 0.005 |
| | HE | 2.069 | 0.02 |
| | HG2 | 2.580 | 0.02 |
| | HG3 | 2.499 | 0.02 |
| | CA | 55.13 | 0.05 |
| | HN | 8.379 | 0.005 |
| | CE | 16.60 | 0.05 |
| **GLY 6** | HN | 8.513 | 0.005 |
| | HA2/HA3 | 3.859 | 0.02 |
| **PHE 7** | HA | 4.575 | 0.005 |
| | HB2 | 3.170 | 0.02 |
| | HB3 | 3.009 | 0.02 |
| | CA | 57.83 | 0.05 |
| | HN | 8.216 | 0.01 |
| | HD1/HD2 | 7.234 | 0.01 |
| | HE1/HE2 | 7.353 | 0.02 |
| | HZ | 7.234 | 0.01 |
| **GLY 8** | HN | 8.465 | 0.005 |
| | HA2/HA3 | 3.879 | 0.02 |
| **LEU 9** | HA | 4.402 | 0.005 |
| | HB2/HB3 | 1.659 | 0.05 |
| | HD1 | 0.950 | 0.02 |
| | HD2 | 0.901 | 0.02 |
| | CD1 | 24.580 | 0.05 |
| | CD2 | 23.270 | 0.05 |
| | HG | 1.659 | 0.05 |
| | CA | 53.83 | 0.05 |
| | HN | 8.132 | 0.005 |

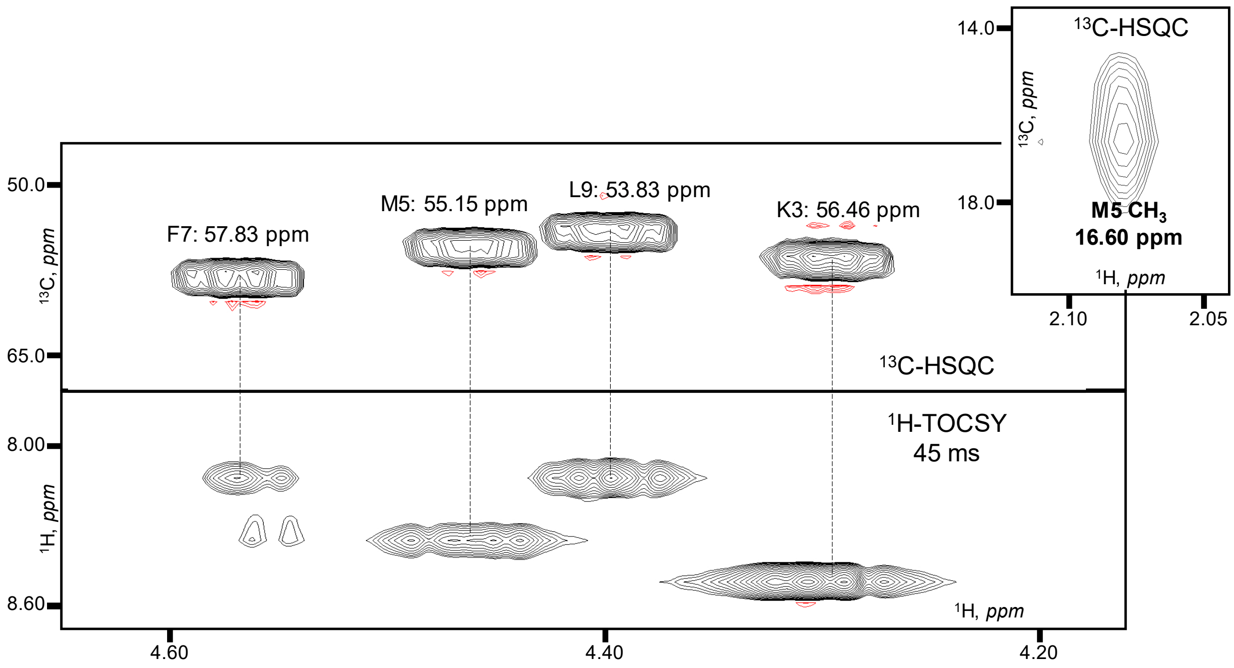


**Figure S1.** Spectral regions for the chemical shift assignments of the $^{13}$Cα for non-glycines residues. The $^{1}$Hα chemical shift values for the $^{1}$Hα-$^{13}$Cα resonances in the $^{13}$C-HSQC and $^{1}$Hα-$^{1}$H$_{N}$ in TOCSY (45 ms) spectra were unambiguously linked. The inset (top-right): Met5 methyl group signal. Other methyl signals (e.g. for L9) are readily identifiable as well.

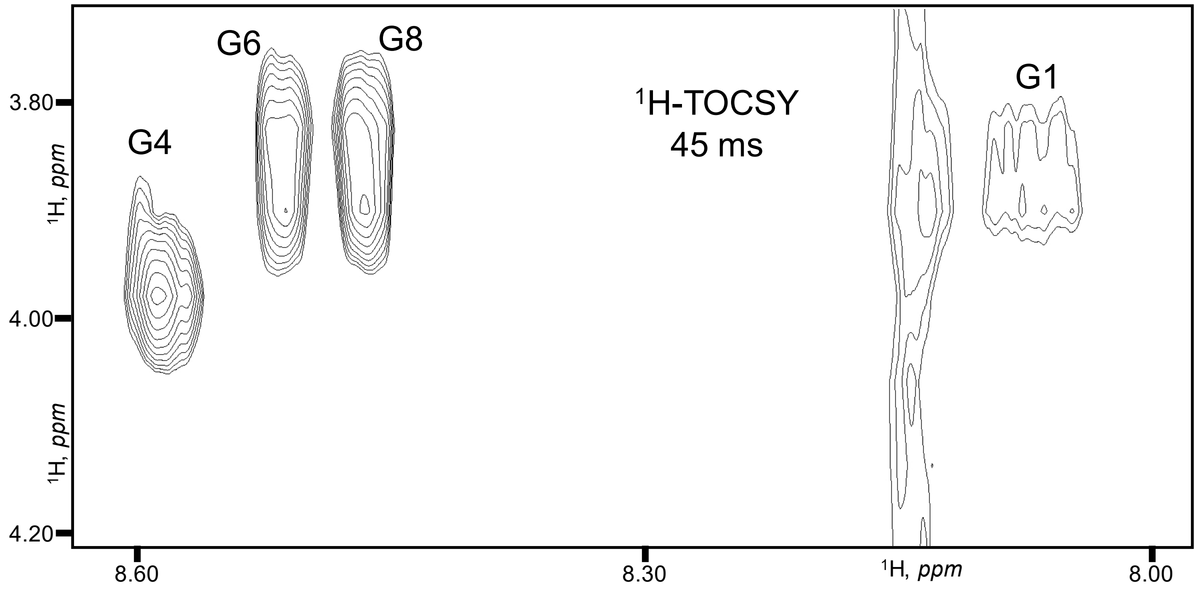


**Figure S2.** The 2D $^{1}$H-TOCSY spectrum, $^{1}$H$_{N}$-$^{1}$Hα region. The G1 cross-peak is the weakest among the four ones labeled.

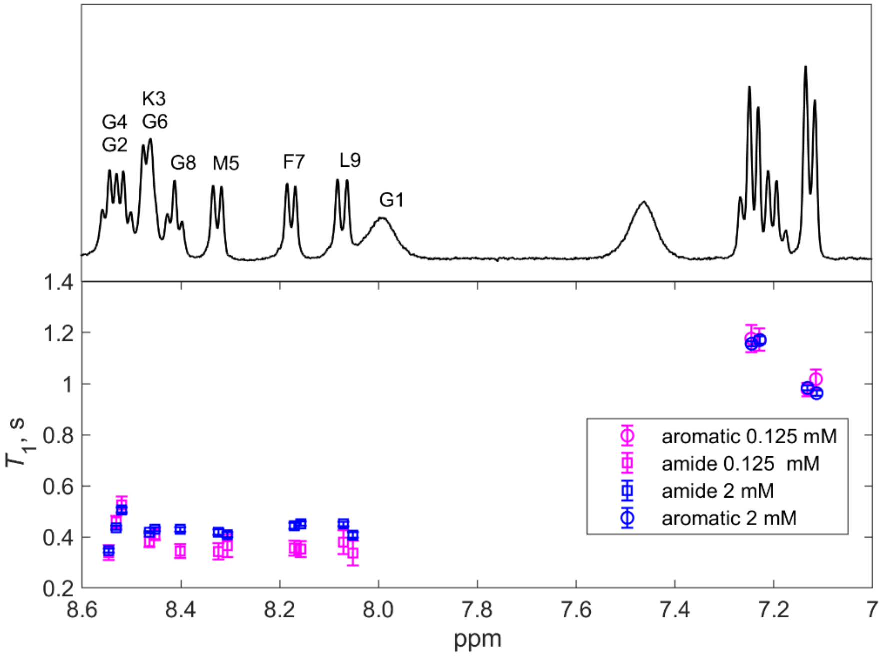


**Figure S3.** Concentration dependence of proton longitudinal NMR relaxation times measured with the inversion recovery technique using the one-dimensional water gate pulse sequence at 9.4 T, 7 °C and pH 2. Top: 1d NMR spectrum at 2 mM concentration with the amide region labeled. Bottom: $T_1$ versus spectral position. Eleven relaxation delays up to 2 s were used in magnetization decay collection. 24 scans were taken for the 0.125 mM sample, and 8 scans for the 2 mM sample. The delays between scans was set to 10 s.

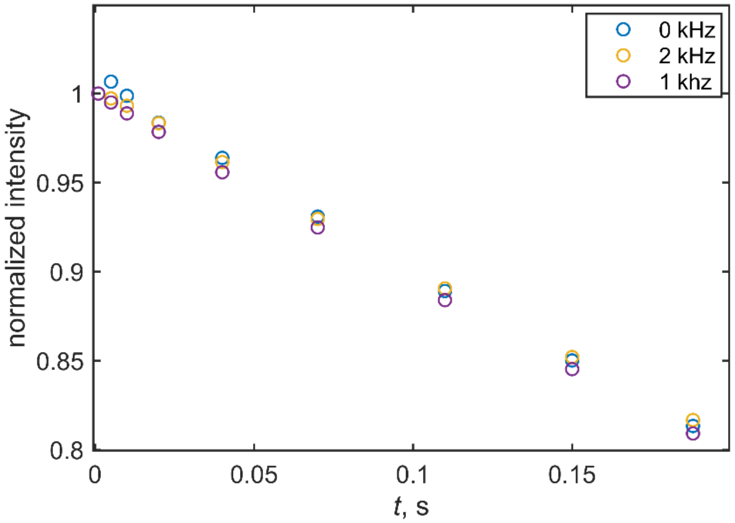


**Figure S4.** Examples of $^{15}$N $R_{1\rho}$ magnetization decay curves for several values of the off-resonance offsets, indicated directly on the legend. The data were collected at 11.4 T, 12 °C, pH 2 and the spin-lock field strength of 2.5 kHz at the selectively labeled amide F7 site. The curves represent the average of the two peaks, which are split by $H_\alpha$ scalar coupling interaction.

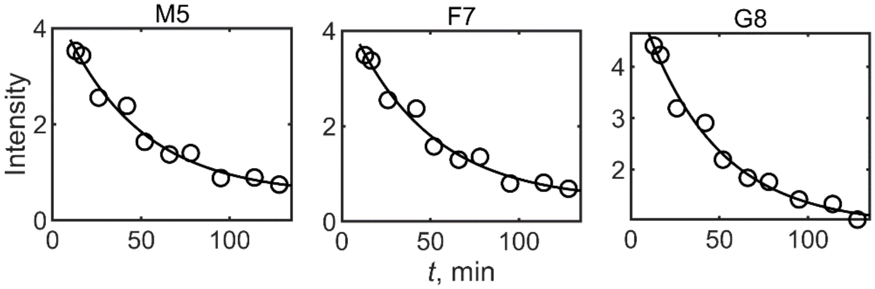


**Figure S5.** Examples of hydrogen-deuterium exchange decay curves fitted to a mono-exponential function with the baseline. Intensities is shown in arbitrary units. The curves represent the average of the two peaks which are split by $H_\alpha$ scalar coupling interaction. The data were collected at 9.4 T, at the sample concentration of 0.9 mM, at 7 $^{o}$C and pH 2. One-dimensional proton NMR spectra were collected with the water-gate solvent-suppression.
